# Prediction of mycotoxin response of DNA-wrapped nanotube sensor with machine learning

**DOI:** 10.1101/2023.09.07.556334

**Authors:** Y. Rabbani, S. Behjati, B. P. Lambert, S. H. Sajjadi, M. Shariaty-Niassar, A. A. Boghossian

## Abstract

DNA-wrapped single-walled carbon nanotubes (DNA-SWCNTs) have demonstrated great versatility as optical sensors. SWCNTs emit a near-infrared fluorescence that is responsive to the slightest changes in their environment, enabling the creation of sensors that can respond to single-molecule fluctuations within the vicinity of their surfaces. The fluorescence response and surface interactions of these sensors are determined by the DNA wrapping sequence. However, the lack of information on the relationship between the DNA sequence and its effect on the SWCNT fluorescence remains a bottleneck for designing sensors specific to analytes of interest. The use of directed evolution was recently demonstrated in order to evolve SWCNT sensors towards mycotoxins through iterative cycles of DNA mutation, screening and selection. In the current work, we use the data acquired during the directed evolution of DNA-SWCNT sensors to train machine learning (ML) algorithms. Artificial neural network (ANN) and support vector machine (SVM) methods were used to predict the response of DNA-SWCNT sensors to the mycotoxin. The reliability of the models was assessed through cross-validation. The cross-validated ANN and SVM models were able to accurately classify the various DNA-SWCNTs as yielding either a high or low fluorescence response with an accuracy of 73 and 81%, respectively. The models were further tested on alternative similar and dissimilar DNA sequences outside of the initial training dataset. The ANN model showed a better ability to predict dissimilar DNA sequences resulting in a high sensor response in comparison with the SVM model. In addition, the possibility to combine the two SVM and ANN models with directed evolution methods was investigated. The experimental results showed that the SVM model was able to predict the response of DNA-SWCNT sensors with 95% accuracy. Finally, the Hierarchy and k-means++ clustering methods were used to examine the similarity and dissimilarity of each DNA sequence at every stage of our investigation. In this work, we show that the application of ML algorithms to directed evolution libraries of DNA allows one to accurately map the performances of DNA-SWCNT sensors within a particular DNA sequence space. Moreover, the computational success of this mapping provides a framework for replacing current empirical approaches with the rational design of DNA sequences for SWCNT sensing.

## 1. Introduction

Nanosensors have unique properties that open up new possibilities for a variety of fields including environmental monitoring, health care, and food industries [1]. Single-walled carbon nanotubes (SWCNTs), which can be represented as graphene sheets that have been wrapped up into seamless cylinders, demonstrate unrivaled physical, mechanical, and chemical capabilities providing an eminent platform for designing the nanosensors [2]. Their near infrared (nIR) emission upon visible light excitation of SWCNTs has attracted particular interest for optical biological applications as this nIR emission spans in the bio-transparency windows [2], [3]. Because SWCNTs can respond to the presence of molecules near their surface with a change in their fluorescence in terms of fluorescence intensity and/or a shift in the emission wavelength, they make good candidates as optical biosensors. In this context, the surface of such SWCNTs sensors can be non-covalently modified with biological molecules, such as DNA and proteins, for a given optical biosensing application [4]. In the case of DNA, single-stranded DNA (ssDNA) interacts with the SWCNT surface in a manner that depends strongly on the DNA sequence, resulting in sequence-specific SWCNT chirality affinity, fluorescence intensity, and analyte selectivity [5], [6]. However, the relationship between DNA sequence and the sensor’s function has yet to be defined [7]. To date, experimental methodologies cannot allow to investigate in-depth such relationship in reasonable time-scales due to the extremely large number of possible DNA sequences, e.g. 4^30^ for a typical ssDNA of 30 nucleotides used in combination with SWCNT [8], [9]. In this regard, the current bottleneck for the development of such optical sensors is the researchers’ ability to imagine, perform and understand the system with a limited number of experiments.

The introduction of machine learning has dramatically driven scientific development by allowing researchers to run experiments in parallel with a variety of learning tools, enabling them to swiftly optimize parameter spaces to reach the desired criteria [10]–[13]. Various studies have been conducted in recent years to examine the behavior of DNA-SWCNT sensors with machine learning. Yang *et al.* (2019) used machine learning methods to investigate the impact of several short patterns in a DNA sequence for SWCNT chirality separation [14]. In 2021, Yaari *et al.* developed a machine learning model for the simultaneous detection of multiple cancer biomarkers such as HE4, CA-125, and YKL-40 in biofluids by training ML with 11 known DNA structures along with 12 SWCNT chiralities [15]. Lin *et al.* (2022) used machine learning to find optimal DNA sequences for sorting individual SWCNT chiralities [16]. Other research studies have used machine learning tools to investigate the influence of DNA sequence on the corona phase molecular recognition [17] and the detection of serotonin [18] using DNA-SWCNTs. All these studies demonstrate the strength of machine learning tools for optimizing the DNA wrapping around SWCNT for various fields of applications. Yet, this growing area of research could also greatly benefit from the exploration of machine learning with other methods.

To date, one of the most efficient ways of finding an ideal DNA sequence for optimizing DNA-SWCNT sensors is directed evolution. This was, for example, used for the engineering of the fluorescent properties of dopamine sensors [8] or to develop multiplexed sensors for a new class of analyte like mycotoxins[19]. However, this procedure limits the search to only a fraction of all possible sensors and therefore yields only partially optimized sensors [20]. In addition, current limitations in screening methods for DNA-SWCNT complexes limit the throughput of this technique. Machine learning, on the other hand, could help to solve the limitations currently encountered for the design of DNA-SWCNT sensors. Machine learning relies on initial information input regarding the sensors. Due to the extremely large optimization space, resulting from the extensive combinatorial DNA sequence possibilities, utilizing appropriate DNA-SWCNT sensors as training input can improve the efficiency of machine learning. This is, for example, the case with data extracted from directed evolution experiments as all the DNA sequences have a close relationship with each other and result in complexes reacting to some extent to the desired analyte.

In this study, various supervised machine learning algorithms have been utilized to anticipate the behavior of a DNA-SWCNT sensor the Fumonisin B1 (FB1) mycotoxin. The cross-validation approach was employed to consider a larger data space of DNA sequences, enabling to improve the reliability of model training. Moreover, the efficiency of several machine learning approaches were investigated for various conceivable DNA sequence spaces. Finally, the models’ prediction results were also employed to design new DNA sequences for experimental testing. This research provides additional insight into the understanding of the relationship between DNA sequence and the properties of the DNA-SWCNT sensors, as well as providing a comparative study on the use of different machine learning algorithms for DNA-SWCNT engineering. The directed evolution technique, with the assistance of an ensemble of two machine learning models, provides a strong tool for the rapid selection of a suitable DNA sequence for a specific analyte detection in future research.

## 2. Result

Machine learning (ML) is a powerful technique that can be used to accelerate the directed evolution method in order to find a DNA-SWCNT complex capable of detecting an analyte selectively. The ML algorithm trained by the directed evolution data predicts the DNA-SWCNT biosensor response to an analyte with a shift in the emission wavelength. The initial dataset for the training of machine learning was obtained from directed evolution experiments[19]. Then, the machine learning model was used to predict new DNA sequences with high shifting responses, which are subsequently selected for the screening step (**Fig. 1**)

**Figure 1.**
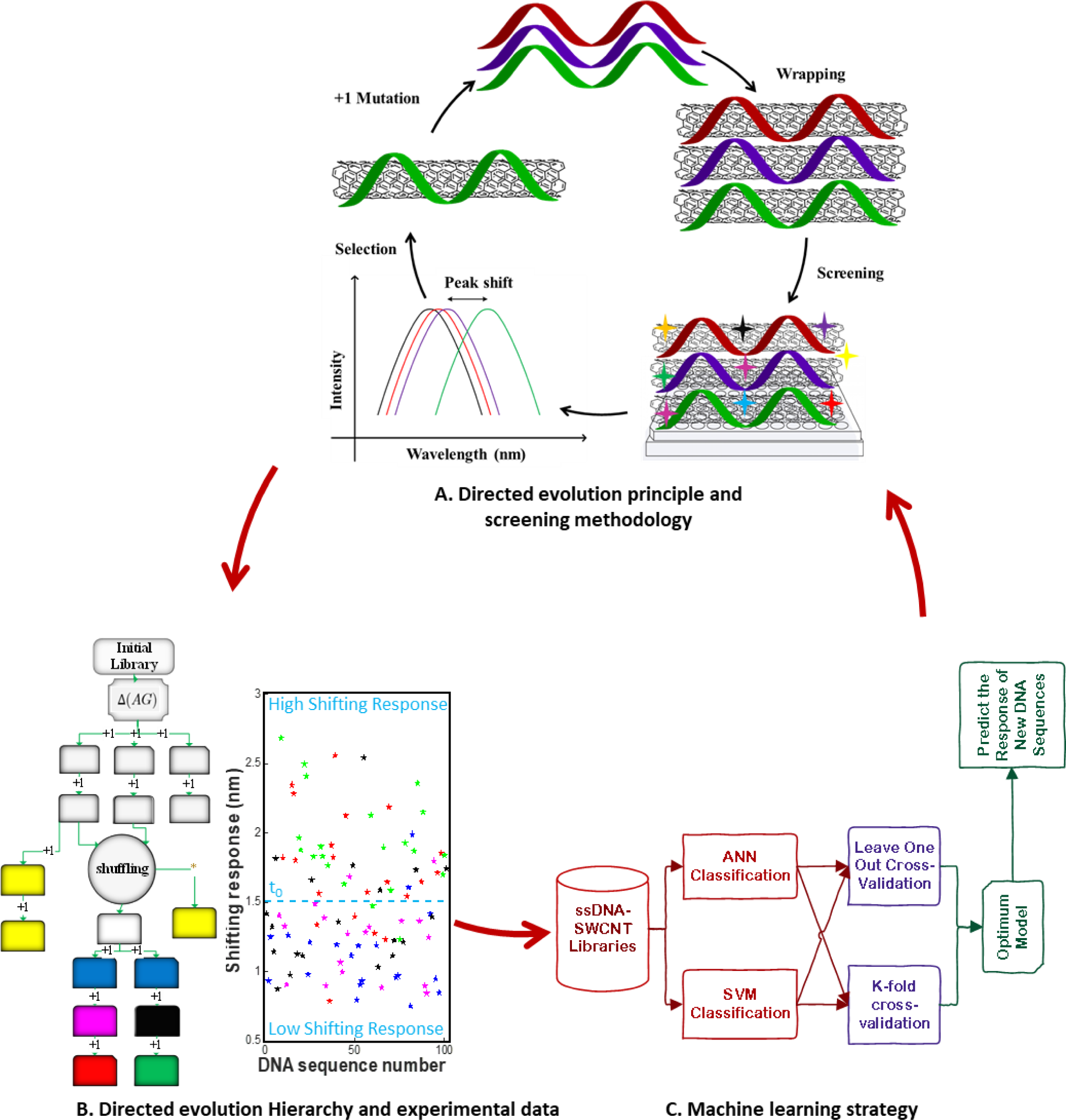
Directed evolution method and Machine learning (ML) procedure for the prediction and enhancement of shifting response for detection of mycotoxin by DNA-SWCNT sensor. A) Directed evolution process for detection of FB1 by DNA-SWCNT sensor. B) Directed evolution hierarchy of DNA sequences mutations and the scatter plot of experimental data: each color in the scatter plot is associated to the same color in the direct evolution hierarchy. The shown threshold (t_0_,blue dotted line) was defined for the classification of data in 2 classes: High shifting response (class 1,top) and low shifting response (class 0,bottom) of DNA-SWCNTs. C) Scheme of how to employ machine learning approach in the directed evolution cycle and the classification procedure of artificial neural network (ANN), and support vector machine (SVM). First ANN and SVM models were trained and evaluated with 101 data sets by cross-validation. Then, the ensemble of the two models was used to predict new DNA sequences that have a high probability of shifting responses, which would be useful for the next screening step of the directed evolution method. * The shuffled sequences were created with a different cutting size (6 nucleotides instead of 5)

As illustrated in **Fig.1A**, an initial screening on a library of diverse DNA-SWCNT complexes yielded the Δ(AG) sensors exhibiting a red-shift for the (7,5) chirality as a response to the addition of the FB1 mycotoxin. Since the response of DNA-SWCNT is a function of DNA sequence, the response was engineered through mutating the initial Δ(AG). A mutation rate of 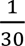 in a 30-bases long DNA sequence results in 90 new DNA sequences with a potential improved response towards FB1 for each round of mutation. In each step, after the introduction of a new random mutation, new DNA-SWCNTs were screened for their response towards FB1 and the sensors with an enhanced response were selected for further mutation. **Fig.1B** shows the directed evolution hierarchy in which after two mutation rounds, another technique of directed evolution called shuffling was utilized to improve the DNA-SWCNT response even further. The data associated to shuffling was chosen to train machine learning as this approach created a more diverse combinations of DNA bases in comparison to the last cycles. The shifting responses of DNA-SWCNTs are plotted in **Fig.1B**. Each color in the scatter plot is associated to the same color in the directed evolution hierarchy. The average value of shifting responses of all DNA-SWCNTs was defined as a threshold (t_0_) allowing classification of data into two classes: low (class 0) and high (class 1) shifting responses.

In this study, artificial neural network (ANN) and support vector machine (SVM) were used as two different supervised machine learning methods to classify data in two separate high and low shifting response classes. First, the entire data was randomly divided into training, validation, and test sets [21]. Then, to make a reliable model, K-fold and Leave-one-out were used as cross-validation methods for the prediction of the next experimental steps (**Fig.1C**). After finding the optimal ensemble of these two models through cross-validation, the new DNA sequences, which have a high probability of shifting response, were predicted for the next step of the experiment.

At first, to investigate a model with the highest precision with respect to different DNA structures in the DNA-SWCNTs sensors for detection of FB1, DNA sequences were converted into a comprehensible language for machine learning methods. One-hot encoding method was implemented for each A, G, T, and C nucleotide resulting in a 4*30 binary matrix as a representation of each DNA sequence, and converted to a feature vector with a size of 120 for machine learning input.

To begin, the models were randomly trained using 70% of the experimental data. To prevent the model from overfitting during training, 15% of the data was used as validation data (**Fig. S1**) and 15% of the data was considered as the final test set [18]. The evaluation parameters have been reported in **Table 1**. The result showed that the model had high accuracy for the test data (86 %). The area under the receiver operating characteristic (ROC), (**Fig.2A**) curve, called AUC, was measured for different datasets. A higher AUC means a better classification performance in most cases; e.g. an AUC value close to 1 corresponds to the ideal classifier. In this study, the AUC was equal to 0.85 for the test data. The results of the ANN classification for test data with a probability of DNA in each class were compared to experimental shifting response values in response to FB1, as shown in **Fig.2B**. Most of the data with a high probability was predicted accurately in either class 1 or 0 with a high or low shifting response, respectively, except for two misclassified DNA sequences. The classification result of training and validation data were shown in **Fig.S2**. Comparing the results of training and validation data showed that the model has a high goodness of fitting without overfitting and underfitting despite the small dataset used. Although the same data distribution was used for the validation and test data, the accuracy of the test data is a bit higher than validation. It may be because the test dataset’s behavior is a little bit easier to predict and closer to the behavior of the training dataset. It is therefore not reliable to use this model to predict similar and distinct sequences altogether. As a result, in the following, the model was trained using several cross-validation techniques to resolve this problem as well as widening the DNA sequence space that can be predicted by the model.

**Figure 2.**
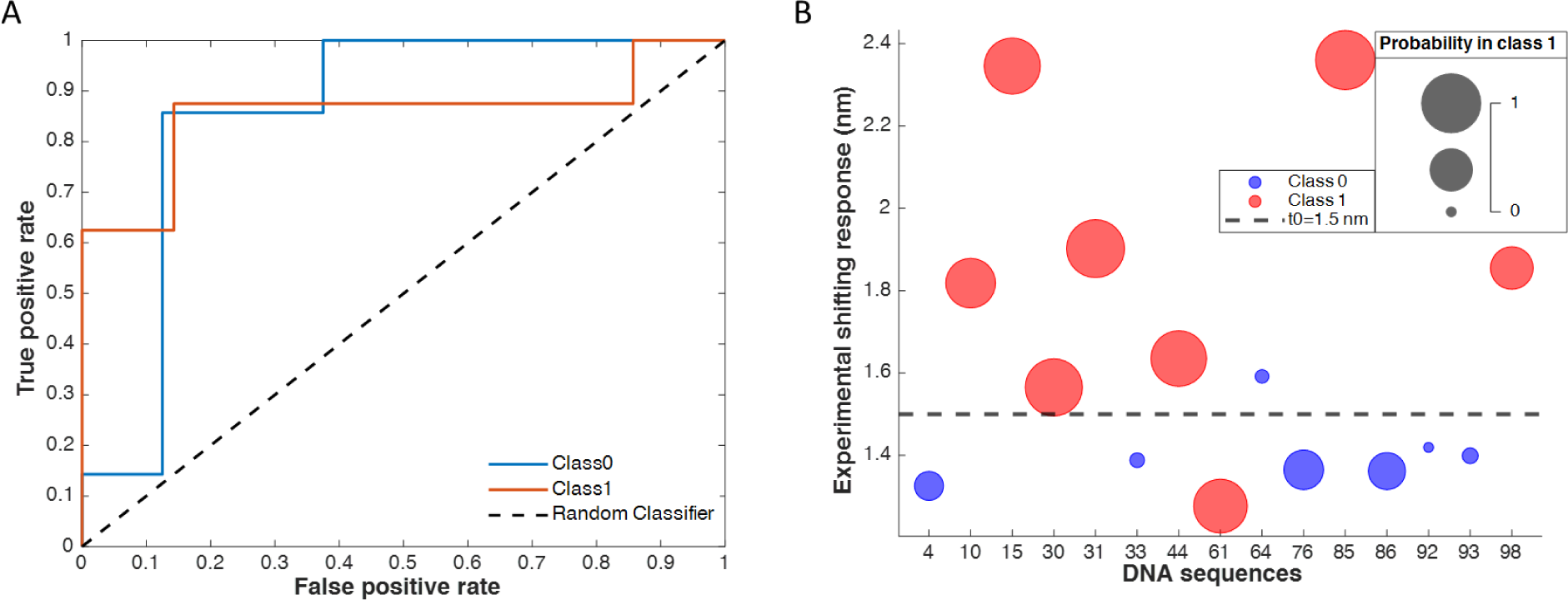
A) The ROC curve for the test data B) Comparing the experimental shifting response values with ML probability classifications of different DNA sequences serves as test data for the detection of the FB1 mycotoxin. The size of circles represent the probability values and the colors represent each class(blue for class 0, red for class 1).

**Table 1.** Evaluation parameters of ANN classification model for training, validation and test data.

|  | Accuracy | AUC | Precision | Recall | True Positive | False Negative | False Positive | True Negative |
| --- | --- | --- | --- | --- | --- | --- | --- | --- |
| Train | 0.80 | 0.88 | 92 | 77 | 20 | 11 | 3 | 37 |
| Validation | 0.79 | 0.80 | 75 | 85 | 6 | 1 | 2 | 6 |
| Test | 0.86 | 0.85 | 85 | 85 | 7 | 1 | 1 | 6 |

### 2.1. Selecting the best classification model with cross-validation

Although the previously discussed model based on ANN was quite accurate in term of prediction, developing an ML dependable model to predict future experiments with high shifting responses necessitates the use of cross-validation procedures that takes into account all DNA-SWCNT system behavior. Therefore, two cross-validation approaches, K-fold and Leave-one-out, applied on two different ML methods, ANN and SVM, have been examined. A comparison of prediction results of these ML methods with the experimental data has been presented in **Fig.3**. The data distribution density in two classes by different ML models has been shown in **Fig.3A**. In the SVM model, the high data density in the first and third quartiles indicates prediction of high and low shifting responses with high probability. The results show that the SVM model performed better since it has classified the data into two distinct classes with high presence probability values and a narrow distribution in each class. The accuracy of SVM and ANN with K-fold cross-validation models was 0.81 and 0.71, respectively. The accuracy of SVM and ANN with Leave-one-out model was also 0.81, and 0.73, respectively (**Table S1**). In **Fig.3B**, the ANN model (coupled with leave-one-out cross-validation) prediction was compared to experimental shifting response values to detect FB1 using various DNA-SWCNT sensors. The majority of the data were almost accurately predicted in the high shifting response classes and DNA sequences with shifting responses larger than 2 nm were correctly classified with high accuracy. **Fig.3C** shows the results of the SVM model prediction using leave-one-out cross-validation. The majority of the data with a high and low shifting response (more than 2 nm and less than 1.2 nm) were predicted with a probability greater than 80% in the correct class with high accuracy. The DNA sequences with a shifting response higher than 2 nm were classified in the right class by all models but the SVM model’s cross-validation results indicated that the model could accurately classify data into two classes for all ranges of shifting response values with a higher probability of prediction. The ROC curve for ANN and SVM models using the Leave-one-out cross-validation procedure is presented in **Fig.3D** and **3E**. AUC for the ANN and SVM models by using Leave-one-out cross validation was 0.75 and 0.78, respectively (**Table S1**). The results obtained from ANN and SVM methods with K-fold cross-validation are compared in **Fig.S3 and S4**. Also, the result of different kernel functions in SVM was reported in **Table S2**. In addition, the effect of changing the threshold (t_0_) on the model’s classification has been studied and is shown in **Fig. S5**. The optimum value of t_0_, determined to be 1.5 nm, was based on the best accuracy value achieved by the ML model. The accuracy was negatively affected by increasing and reducing the threshold from the average value (t_0_=1.5nm). It should be noted that this parameter completely depends on the shifting response distribution, and the average value in this study had a high level of accuracy and data classification. Although the comparison with other research papers may not be entirely precise due to the use of different libraries of DNA sequences, the examination of accuracy in relation to similar studies revealed a significant impact on model accuracy when utilizing the directed evolution library as input data for machine learning (ML) training, resulting in an increase of up to 0.8 [14], [22]. For instance, Kelich *et al*. [14], [22] employed a DNA library derived from SELEX to explore DNA-carbon nanotube sensors for serotonin, achieving an accuracy range of 0.64 to 0.71. Additionally, Yang *et al*. [14], [22] reported a precision range of 0.48 to 0.6 when learning to predict single-wall carbon nanotube-recognition DNA sequences.

**Figure 3.**
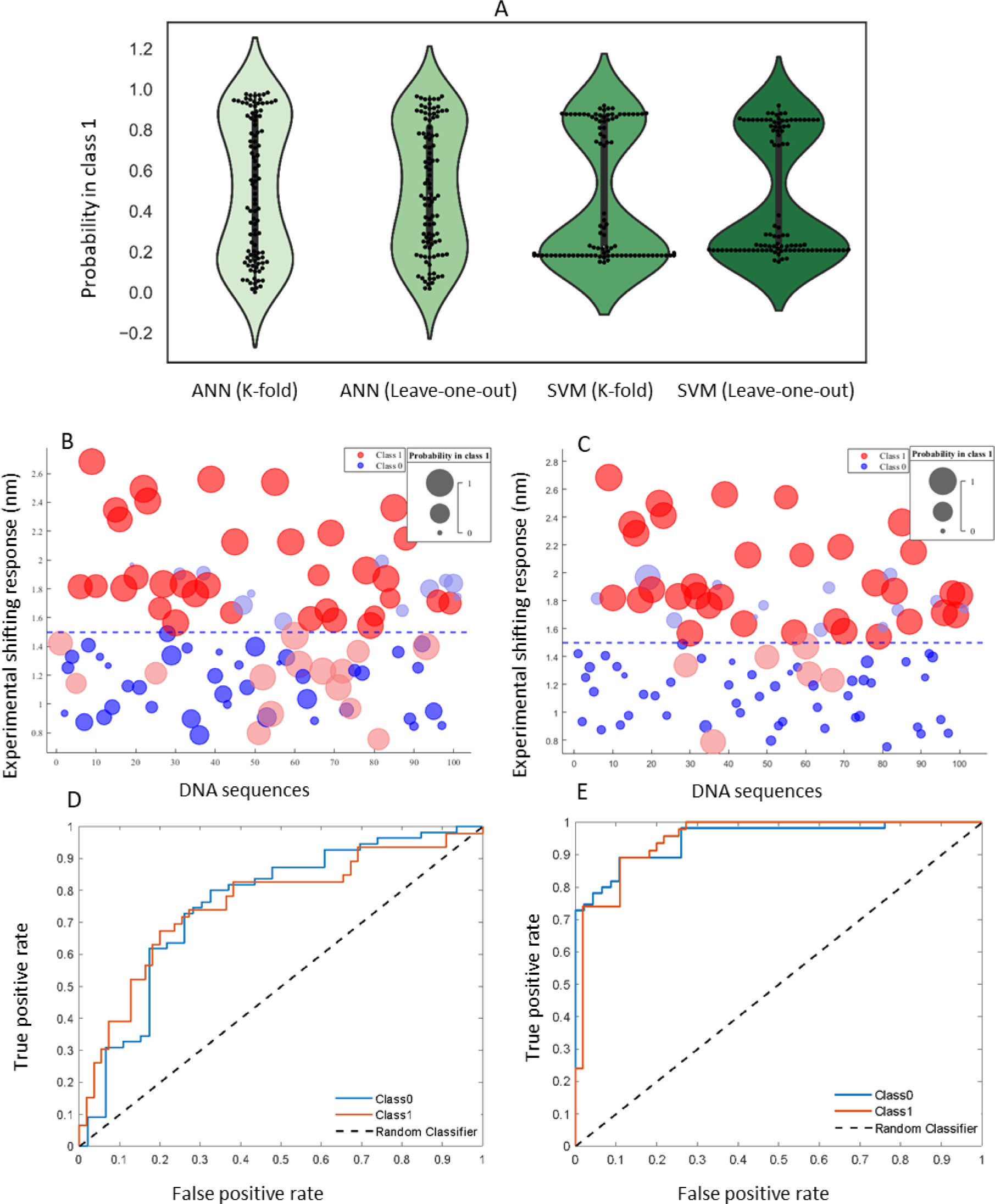
The comparing of ANN and SVM models with experimental shifting response: A) The distribution density of the prediction results in two class for all ML models. The DNA sequence classification and presence probability of each DNA sequence in each class predicted by the ANN (B) and SVM (C) model with Leave-one-out cross-validation compared with experimental shifting response values to detect FB1 by different DNA-SWCNT sensors. The size of the circles shows the probability values. Low shifting responses (<1.5) show class 0 with blue color, and high shifting responses (≥1.5) show class 1 with red color, respectively. In addition, the dark color indicates correct class predication and the light color shows the misclassification. The ROC curve for the ANN model (D) and SVM model (E) with leave-one-out cross-validation (AUC = 0.73 and 0.78).

### 2.2. Model’s ability to predict new dissimilar DNA Sequence

In this study, we trained machine learning algorithms by SVM and ANN methods based on data gained from directed evolution (3 last steps of mutation) which are in a narrower range of DNA base combinations space and high similarity among them. However, as a huge number of potential DNA base combinations with nucleotide lengths of 30 is possible, we need also to evaluate the ability of the model to predict the behavior of alternative DNA sequences in various regions of combinations and less similarity to the training dataset. As a result, the model’s potential to predict alternative DNA sequences in various regions was investigated. Established models based on SVM and ANN were used to predict the classification outcome of the new data gathered from both mutants created from DNA shuffling and from individual non-shuffling mutations, as shown on the directed evolution hierarchy (**Fig.1B**, yellow boxes), and presented in **Fig.4**. **Fig.4A** presents the proportions of data correctly or incorrectly classified into either class 1 or 0 by both the ANN and SVM models. It is evident from the graphs that the majority of the data which has a high experimental shifting response (>1.5nm) was accurately assigned to class 1. For more details, the results of the confusion matrix (COF) and ROC curves obtained from the prediction model are shown in **Fig. S6**. In addition, **Fig.4B** provides a comparative analysis between the model’s predictive outcome (Probability in class 1) and the experimental shifting response (shifting response higher and lower than 1.5 nm as class 1 and 0). The results revealed that ML classification using various models was unable to effectively classify data in class 0, but was a good classifier for class 1. In terms of classifying data from diverse domains, as demonstrated in the directed evolution hierarchy involving dissimilar DNA sequences, the ANN model exhibited superior classification performance compared to the SVM model. Generally, the ANN has the ability to learn and capture complex patterns rather than SVM Model for non-linear mapping. As a result, the SVM model, which is based on the support vector guidance for each feature, is a good predictor for data with similar behavior, but the ANN, which is based on weight adjustment for each feature, is a good predictor for data with complicated dissimilar features [23], [24].

**Figure 4.**
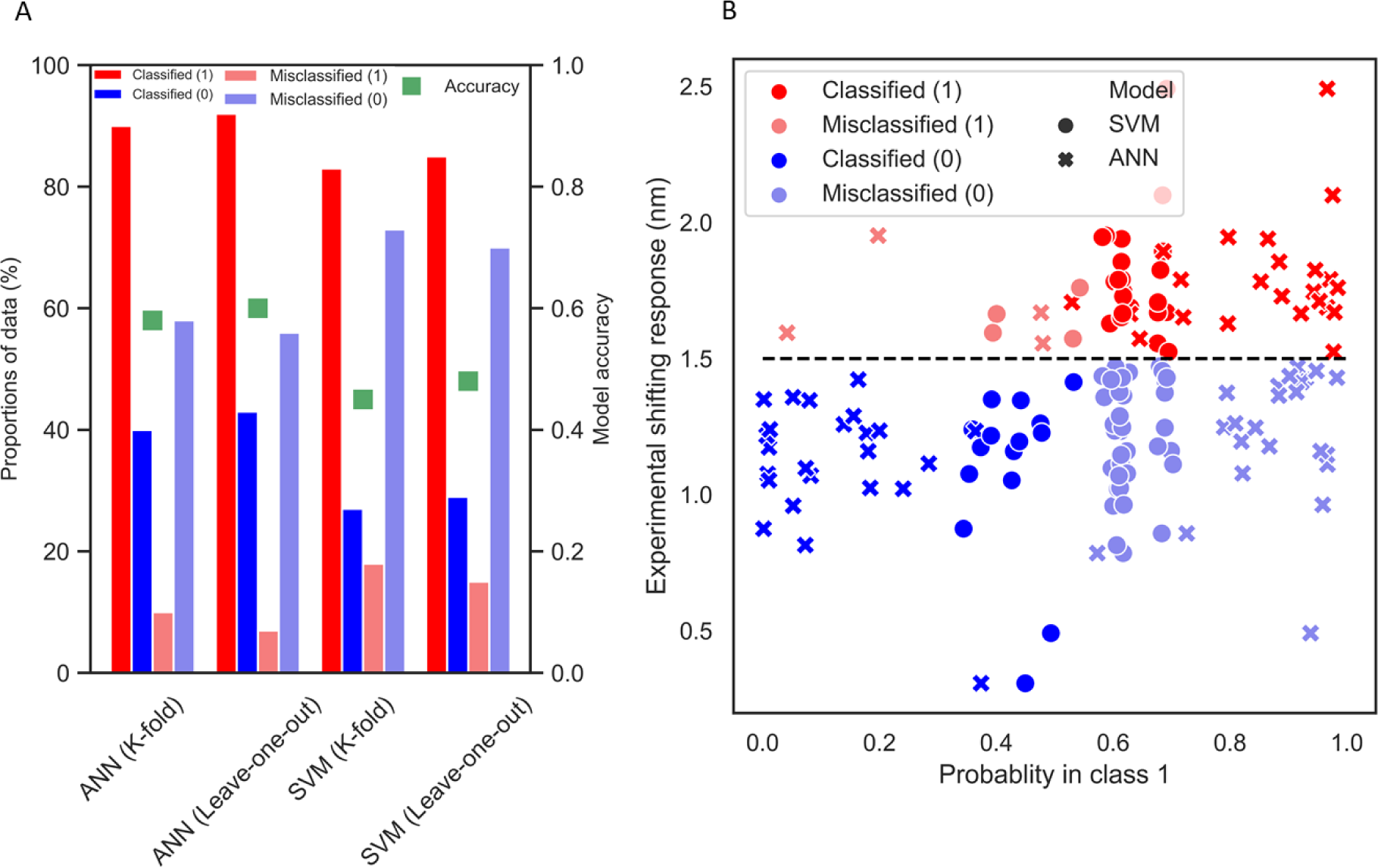
The ability of ANN and SVM models to predict new, dissimilar DNA from distinct directed evolution cycle. The data was selected from the shuffling stage and non-shuffling mutation part combined together as shown in the directed evolution hierarchy (Fig.1B, yellow boxes) and classified by ANN and SVM method. A) Bar plot of the percentage of data in the right class and misclassification with model accuracy for SVM and ANN models. B) Comparison of the result of ANN and SVM with Leave-one-out classification and experimental shifting response. The low shifting response (< 1.5) indicates class 0 and the high shifting response (≥1.5) shows class 1. The dark color indicates correct class predication and the light color shows the misclassification.

### 2.3. The ability of the ML for finding new DNA Sequence in directed evolution method

As described in the acquisition section, with one mutation in a 30-length sequence, 90 DNA sequences were produced in each round of the directed evolution cycle. Following the mutation, all DNA-SWCNTs need to be screened in order to find the optimal mutant, which can be time and resource-consuming. ML can help directed evolution in finding the desired DNA sequence quicker in this stage. As SVM worked better for similar DNA sequences and ANN was more accurate for dissimilar ones, we used ensemble of these two models with Leave-one-out cross validation to predict responses of 90 altered sequences mutated from the best final sequence of directed evolution for the next experimental stage (**Fig.S7**). From 90 mutated DNAs mutation, 22 sequences were predicted with high probability in class 1 (in both models) and were then subsequently evaluated experimentally to detect FB1 (**Fig.5A)**. The experimental outcome of ML predicted sequences is displayed in **Fig. 5B** and compared to ML classification. In **Fig.5C**, the outcome of ML prediction versus the last cycle of directed evolution (black into the green box in **Fig.1B**) revealed that ML was able to find more DNA sequences with higher response rather than directed evolution. Therefore, directed evolution assisted by ML helped to increase not only the number of sequences with improved response increase from 5% to 40% in comparison to conventional directed evolution, but also allowed to increase the absolute response.

**Figure 5.**
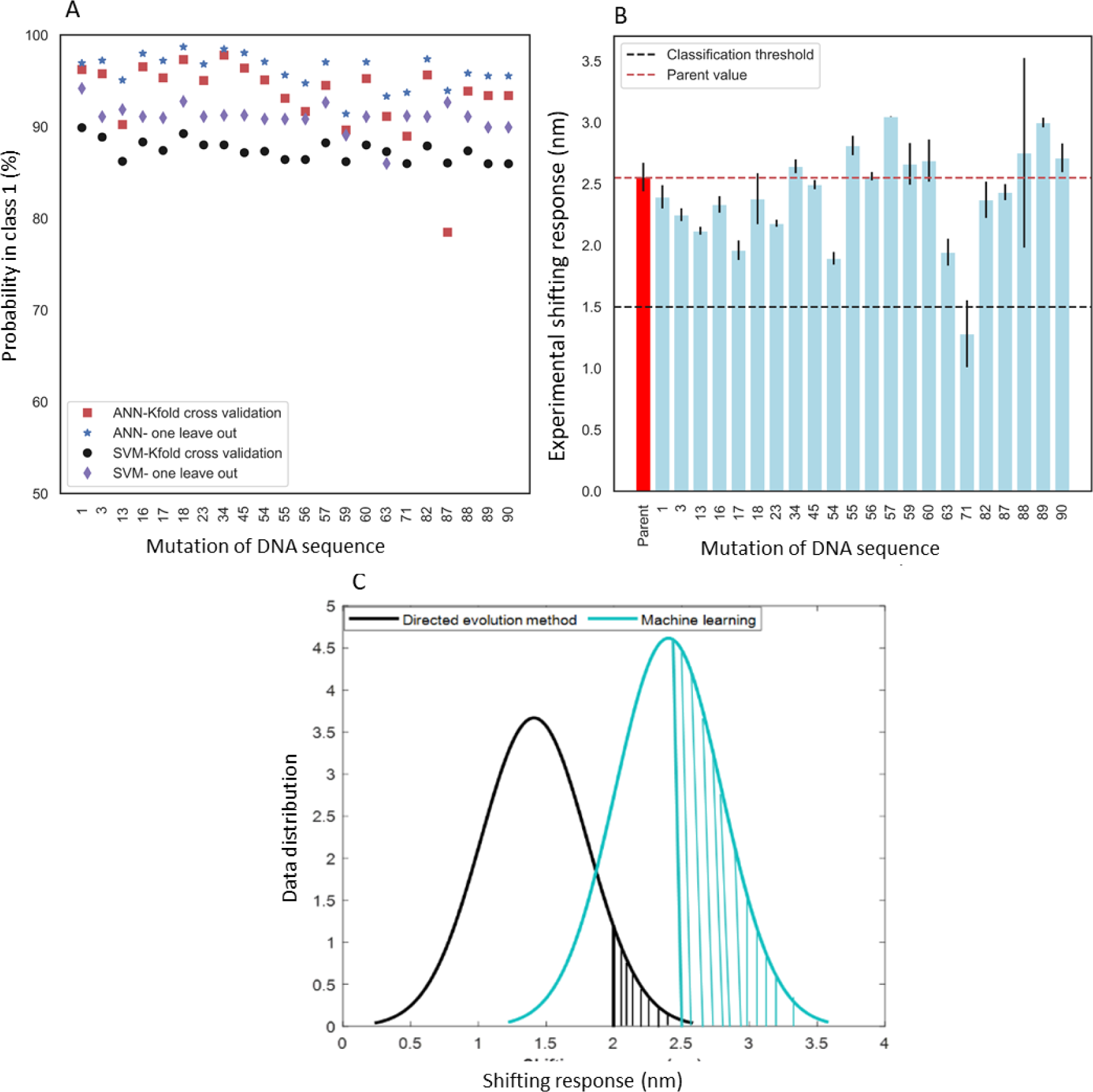
The result of the ML prediction for new mutations compared with experimental data A) Result of ML prediction of DNA mutations that had the highest probability in class 1: the DNA sequences were selected based on the best result of SVM and ANN with Leave-one-out cross validation methods B) Experimental peak shifting results of selected DNAs: sequences were predicted in the right class except for one misclassification(sequence no. 71) C) The result of ML prediction was compared with the previous step of the direct evolution cycle (from the blue to the black box in the hierarchy in Fig.1B)

## 3. Discussion and Conclusion

One of the most efficient ways to find an ideal DNA sequence for increasing the fluorescence response in ssDNA-SWCNTs sensor is the directed evolution method. By narrowing down the unlimited space of DNAs, this technique brings the opportunity to engineer different properties to get the desired biosensor. This procedure is time-consuming although it is an effective way, and machine learning (ML) can be used to boost this method. In this work, a data set of the high-throughput screening of directed evolution method has been used to train ML algorithms. In this regard, artificial neural network (ANN) and support vector machine (SVM) models with K-fold and Leave-one-out cross-validation methods have been used to predict the behavior of DNA-SWCNT sensors for detecting the FB1 mycotoxin. The ensemble of two ML models was used to predict the DNA mutation in combination with directed evolution for generating new experimental screening. In addition, the model’s ability to predict dissimilar DNA sequences based on directed evolution hierarchy was studied for different libraries of DNA sequences. The ability of the ML models to predict similar DNA (produced by mutation in the directed evolution method) and dissimilar DNA sequences has been shown in **Fig.6**. As can be seen at the top of the figure, all DNA sequences have been divided into four groups, each denoted by a different shape. Group A comprises a library of data used for training, Group B contains mutation libraries, and Groups C and D are two different libraries. The efficacy of machine learning classification for these different groups is shown by green face edge colors for correct classification, and red face edge colors for misclassification in both high shifting response (class 1) and low shifting response (class 0). The specified ML models proved to be effective classifiers for the mutation libraries.

**Figure 6.**
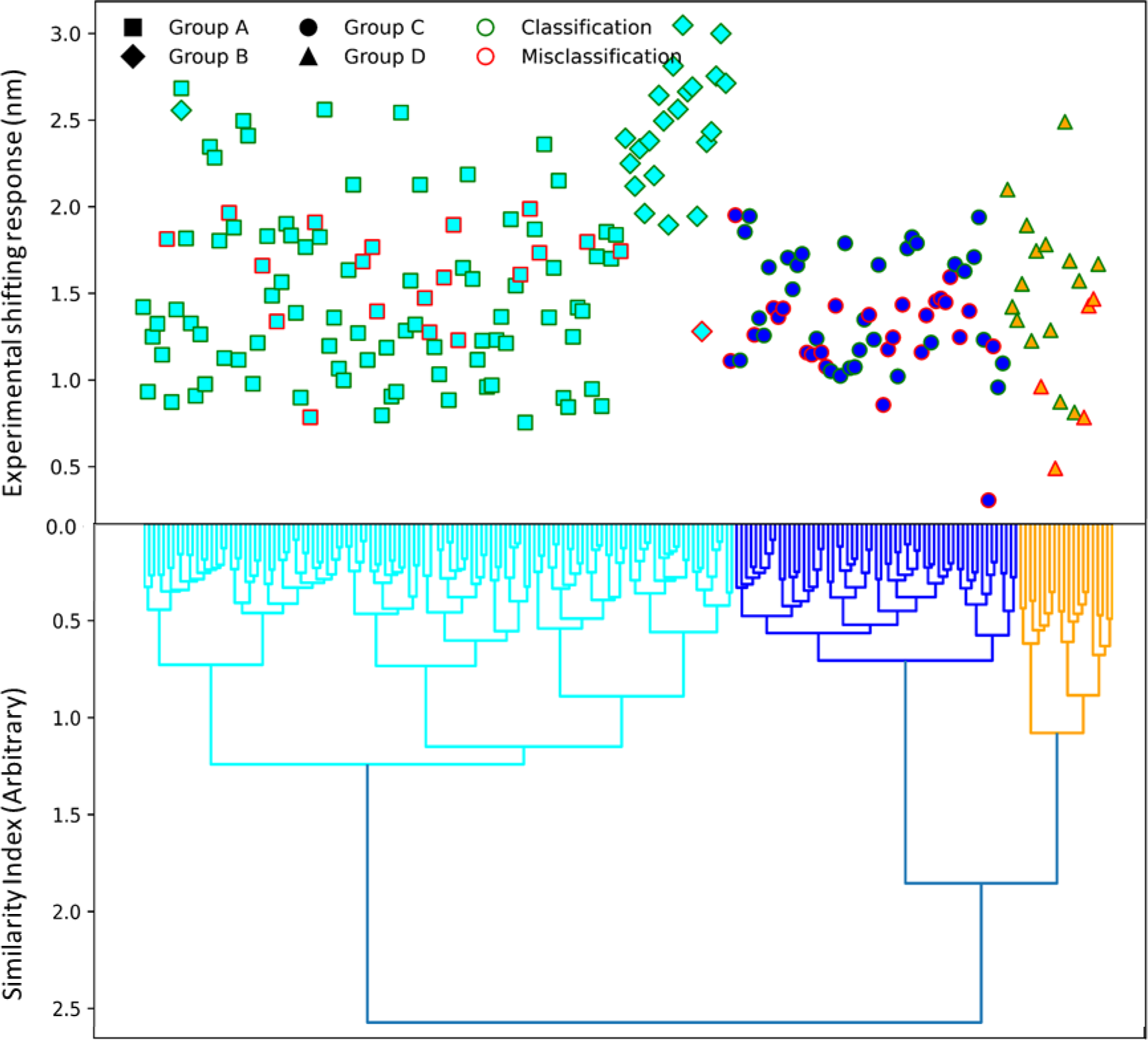
The ability of the ML models to predict similar DNA (produced by mutation in directed evolution method) and dissimilar DNA sequences libraries with comparation of experimental shifting response. Group A is a library of data used for training, Group B is mutation libraries, and Groups C and D are two different libraries. The green face edge colors represent correct classifications, while the red face edge colors indicate misclassifications in both high shifting response (class 1) and low shifting response (class 0). The similarity of different DNA sequences was investigated by hierarchy clustering unsupervised machine learning method. In this dendrogram, distinct clusters are represented by different colors, where each node signifies a unique DNA sequence. The connections or branches between these nodes reflect their similarity. Moreover, the height of these branches indicates the level of dissimilarity or distance between the DNA sequences, with a higher branch suggesting less similarity.

Hence, for better understanding of ML prediction results, the similarity of all DNA sequences was assessed by unsupervised clustering machine learning methods like k-means^++^ and Hierarchical clustering methods [25], [26]. The similarity of the DNA sequences in mutation part (Group B) is close to the DNAs in the training data set (Group A), which is the reason of high accuracy in the mutation part prediction proved experimentally in combination with directed evolution method. In Group C and D, the behavior of DNA sequence is different from the training data set that caused the less accuracy especially in low shifting response rather than the mutation part (Group B). Still, ML model has a goof ability to classify the DNAs with high shifting response.

ML models have an excellent ability to be combined with directed evolution approach because of the huge DNA space. The combination of directed evolution and machine learning can then be utilized to detect distinct analytes as a viable strategy for sensor fabrication engineering.

## 4. Method

### 4.1. DNA-SWCNT data preprocessing

Experimental data provided by directed evolution method was utilized for training of several supervised machine learning approaches. For ML training, shifting responses of (7,5) chirality of DNA-SWCNTs to the FB1 analyte were considered in two classes (High and low shifting response). Each DNA sequence was likewise encoded using the one-hot encoding approach [27], [28] and then transformed to a numeric array as follows:

**Figure 8.**
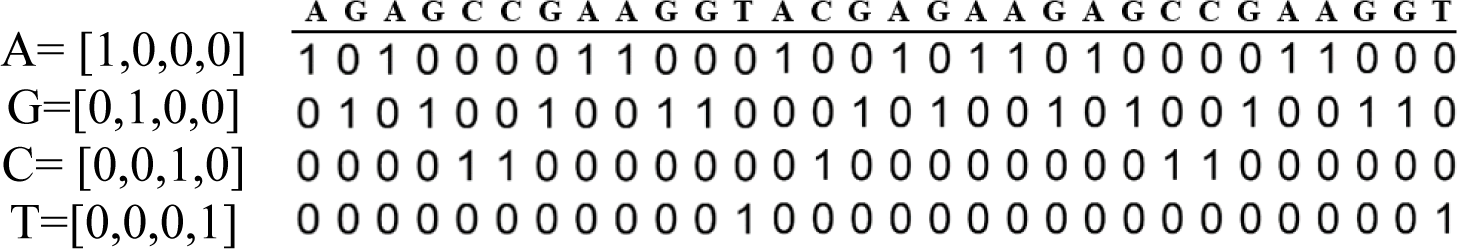
An example of encoding a random DNA sequence matrix with 30 nucleotides by one-hot encoding method.

Then, this matrix (4*30) was converted to the (1*120) array as an input data for machine learning methods.

### 4.2. ANN method

Artificial Neural Network is made up of basic operational pieces that process the network’s incoming data in a parallel way. The inputs multiplied by the modifying weights and bias are summed and then sent via a function to generate the output for that neuron, as indicated in the equation below [29], [30].

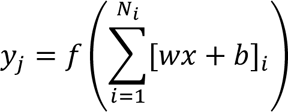

Where *x* and *y* are the *i-th* and *j-th* input and output, respectively, *N_i_* denotes the number of inputs, *w* is the connection weights, *b* represents the bias, and *f* is the operating function of a neural network. For classification, a feed-forward fully connected neural network was utilized which can perform non-linear mapping with variable precision depending on the number of used layers and neurons. Different algorithms were used in the training step, such as Gradient Descent, Gradient Descent with Momentum, Conjugate Gradient, Quasi-Newton and Levenberg-Marquat algorithm [31]. The network was scored using the Softmax algorithm, which divided the results into two classes. Various neural network topologies, such as hidden layers and the number of neurons, were also investigated in order to develop the best model with the least amount of error.

### 4.3. SVM Method

Support vector machine (SVM) is a monitoring-based learning system that is utilized for classification and regression. The SVM has been used to tackle anticipating and optimization challenges. SVM estimates a function linked to dependent (*yi*) and independent variables(*xi*) as *y = f (x),* that is comparable to the equation below:

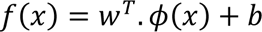

The purpose of this equation is to develop a function with weights (*w*) vector, bias(*b*) factor, and kernel function ϕ(*x*). This may be accomplished by using a collection of data to train an SVM model, using the model’s operating premise previously stated in a prior study [20]. In the SVM model, the features of w and b are computed using the Karush-Kuhn-Tucker theory conditions, where w is defined as in the equation below.

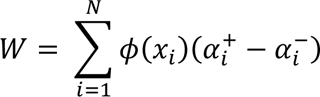

Where, ϕ(*x*) is the core function and *α* is Lagrange coefficient. Since calculating ϕ(*x*) is difficult, a Kernel function is defined in this equation to solve it:

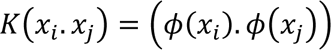

For a feed backward vector classification model, different kernels are employed, including linear, polynomial, quadratic and Gaussian. The model parameters, as well as the kernel parameter, must be carefully computed to generate an effective model for SVM technique.

### 4.4. Cross-validation

Cross-validation was used to increase the generalization and independence of the ML model from training data set. Distinct K-fold and Leave-one-out cross-validation for different ML algorithms were employed for this purpose. The original sample is randomly split into sub-samples of size k in K-fold cross-validation. A single sub-sample is kept as validation data for model testing, and k-1 sub-samples are utilized as training data in the k-subsamples. The cross-validation technique is then utilized exactly once as validation data for each of the k samples after being repeated k times. The data were then averaged to provide a single estimate. This approach has the benefit of repeating random sampling in a way that all observations are used in both training and validation steps. The loss function, J(w), was developed for this technique based on neural network model [32]:

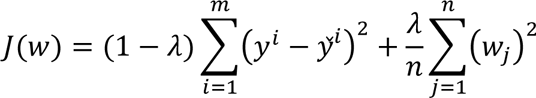

where *Y* and *Ŷ* and are the *i-th* measured and estimated outputs, *wj* is the *j-th* weight, m, and n are the network’s quantity of training data and weights, and λ is the regularization optimization parameter between 0 and 1. A term containing the total amount of squared weights (Right side in Eq.) is added to the *J(w).* To reduce the loss function, the network weights and λ regulation parameters were optimized. The model is fitted to experimental data points by adjusting the weights to minimize the first term of the equation. When the number of k-subsamples is same as all data, K-fold cross-validation is equal to Leave-one-out approach. In other words, only one sample is kept as validation data for model checking in each cycle, while the rest of the samples are used as training data.

### 4.5. ML evaluation parameters

A different variable, confusion matrix (COF), was defined for the evaluation of different machine learning classification models. The matrix compares the actual goal values (experimental data) to what machine learning model predicts. A good model has high TP (true positive) and TN (true negative) rates but low FP and FN rates.

**Figure 10.**
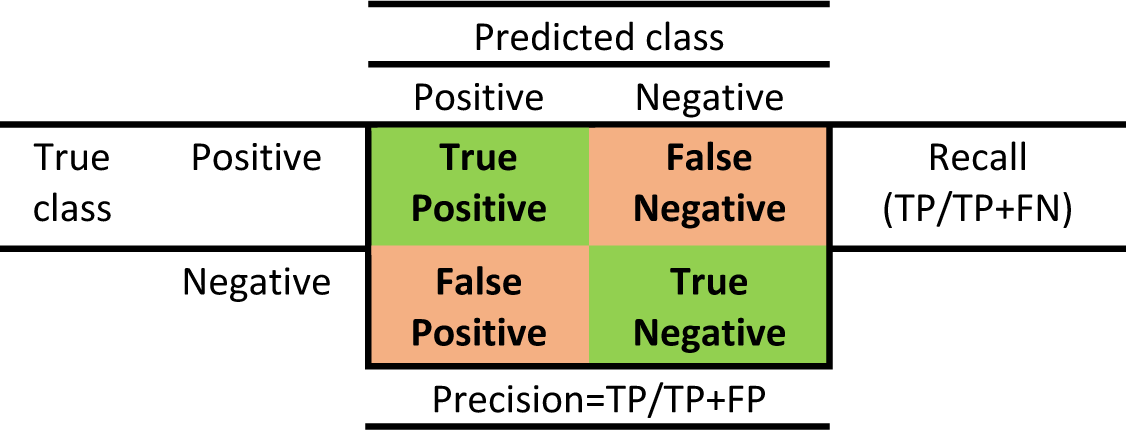
COF matrix definition

True Positive (TP) denotes that the model predicted yes and the actual value confirmed that prediction. True Negative (TN) means the model predicted No and the true or actual value was likewise No. The term false positive (FP) refers to when the model predicted Yes but the actual value was No. The model predicted no, but the actual value was Yes, according to False Negative (FN). The following are the definitions for true positive rate, true negative rate, and accuracy:

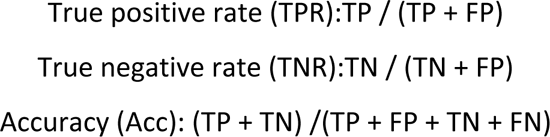

The Receiver Operator Characteristic (ROC) plot is a graphical representation of binary classifier diagnostic performance. The true positive rate is plotted against the false-positive rate to form a ROC curve. ROC curves are advantageous for comparison among several classifiers by combining their performance into a single metric. The area under the ROC curve, abbreviated as AUC, with a 0 to 1 range is a commonly used method serving as a general metric of prediction precision [33]. It’s the likelihood of a randomly chosen positive case being rated higher than a randomly chosen negative case, or the Wilcoxon rank-sum statistic for two samples.

### 4.6. Clustering method for DNA similarity

The machine learning clustering algorithm can group sequences with similar properties and investigate novel sequences from known functions and structures. Clustering DNA sequences is important to see the similarity and dissimilarity of DNA sequences. Each cluster is unique and cluster analysis groups similar data together for further analysis. For the encoding of the DNA structure for clustering, One-Hot encoding method same as previous procedure was used and clustering of the unlabeled DNA data can be performed with the k-means^++^ and hierarchical clustering.

#### K-means method

The K-means algorithm is a non-hierarchical clustering method that initially determines the number of clusters. It works by partitioning samples into groups with equal variance and minimizing the inertia or within-cluster sum-of-squares. A set of N samples is divided into K disjoint clusters via the k-means algorithm, and each cluster’s mean μ_*j*_ is used to describe its samples. The means are often referred to as the cluster “centroids,” however despite existing in the same area, they are rarely actual points from X. The within-cluster sum-of-squares criterion, or inertia, is the goal of the k-means method, which seeks to select centroids that minimize it:

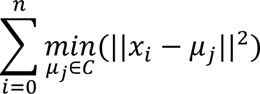

K-means always converge, but to a local minimum. This depends on centroids’ initialization. The algorithm is commonly repeated with different centroids. k-means^++^ initialization helps address this issue.

#### Hierarchical clustering method

A wide family of clustering algorithms known as “hierarchical clustering” creates nested clusters by gradually merging or breaking clusters. A tree is used to depict the clusters’ hierarchical structure (or dendrogram). The unique cluster at the tree’s base has all of the samples, while the clusters at the tree’s leaves each contain a single sample. Each observation begins in its own cluster, which is then gradually combined by the Agglomerative Clustering object to create a hierarchical clustering. The metric employed for the merging approach is determined by the linkage criterion. The following equation-based Euclidean distance measure was employed in this study:

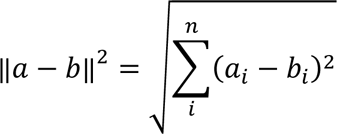

### 4.7. Experimental procedure for prediction of the mutation part with ML

#### ❖ DNA-SWCNTs preparation

DNA-SWCNTs are prepared based on the method described in the previous paper [8]. Briefly, as a first step, 45 mg of super purified HiPCO SWCNT (NanoIntegris, batch HS37-007) was dispersed in 45 mL of 2% (w/w) sodium cholate dissolved in deionized water (2% SC) using homogenizer (PT 1300D, Ploytron) for 20 minutes at 500 rpm and followed by sonication with probe-tip ultrasonicator (1/4 in. tip, Q700 Sonicator, Qsonica) at amplitude of 10% in an ice bath for 1 hour. Afterwards, the suspension was centrifuged at 15000 x g for 10 minutes at 25 °C (Beckmen Allegra). For complete removal of aggregates, Optima XPN-80, Beckman Coulter centrifuge was utilized for 4 hours at 25°C and 164000 x g and 80% of the supernatant was collected. The final solution was concentrated with 3K Amicon ultra filter devices at 4000x g for 30 min and following centrifuge (Beckmen Allegra) was done to remove any remaining aggregates. The absorption of 2% SC-SWCNT solution was measured at 632 nm in a quartz cuvette with Shimadzu UV-3600 Plus spectrophotometer. The solution concentration was calculated with extinction coefficient of 0.036 L.cm^-1^.mg^-1^ and adjusted at 108 mg.L^-1^.

ssDNA sequences (Microsynth, Switzerland) were dissolved in deionized water and the concentration was adjusted to 50 µM measured with Nanodrop 2000, Thermo Scientific. 60 µl of 2% SC-SWCNT (108 mg.L^-1^), 60 µL of DNA solution and 180 µL of methanol (VWR chemicals) were mix in 1.5 ml Eppendorf tube and incubated for 2 hours for DNA and SC replacement. To remove methanol and SC from DNA-SWCNT suspension, DNA-SWCNT was precipitated by adding 866 µL of 100% ethanol (Fisher Scientific) stored at -20°C and 46 µL of 1.5 M NaCl (Sigma-Aldrich) solution in deionized water and stored at -20° for 1 hour and centrifuged at 21130 x g for 30 minutes (5424 R, Eppendorf) to form a pellet from the precipitants. In the next step, the supernatant was discarded and the pellet was washed and vortexed with addition of 1 mL 70% (v/v) ethanol. The process followed by centrifuging the suspension at 21130 x g for 1 hour. The supernatant was discarded and the pellet was dried under a fume hood for 12 minutes. The DNA-SWCNT pellet was resuspended in 300 µL of 100 mM NaCl solution and centrifuged for 30 min and 21130 x g to remove aggregates. The absorption of the DNA-SWCNT suspension was measured by plate reader (Varioskan LUX, Thermo Scientific) at 632 nm using 90 µL of the solution in a 384-well plate. The samples were diluted by addition of 100 mM NaCl solution to adjust the absorption at 0.1.

#### ❖ Screening for mycotoxin FB1

Mycotoxin FB1 (Enzo Life Sciences) was dissolved in dimethyl sulfoxide (DMSO) in a nitrogen-controlled atmosphere glovebox (E-line, GS Glovebox). For Screening, 49.5 µL of each DNA-SWCNT solution was added to a 384-well plate. Then 0.5 ul of FB1 solution (1mM) was added to the wells. As a reference to compare the PL changes, 0.5 µl of DMSO was added as a blank to the same sequence in another well. After 80 minutes incubation time at room temperature, the PL of the ssDNA-SWCNT was recorded and the measurement was done again after 200 minutes of FB1 and DMSO additions. All experiments were done with 3 replicates.

#### ❖ Photoluminescence measurement setup

As indicated in our previous work, a custom-built optical set up was used for recording PL including supercontinuum laser source with a tunable filter unit (SuperK Extreme EXR-15 and SuperK Varia, NKT Photonics). The nIR light reflections were removed by a short-pass filter (890 nm blocking edge BrightLine, Semrock). The excitation light was directed into an objective (M Plan Apo NIR, NA 0.4 air, Mitutoyo Corporation) with 20x magnification. The final illumination spot size on sample was 350 x 350 µm. The emitting light from the sample was collected and directed to an IsoPlane SCT-320 spectrometer (Princeton Instruments). Then, the collected light was dispersed by a grating of 70 lines mm^-1^ and directed to an InGaAs nIR camera (NIRvana 640 ST, Princeton Instruments). Excitation wavelength of laser was 660 nm to monitor (7,5) and (7,6) chiralities. The emission was collected for the wavelength of 900 nm to 1400 nm. PL spectrum analysis was done with MATLAB and the difference in the peak position of (7,5) chirality in FB1 and DMSO was considered as a peak shifting response.

## Acknowledgements

The authors express their gratitude for the funding support received from various sources, including the Swiss National Science Foundation (SNSF), the European Research Council (ERC) through the European Union’s Horizon 2020 research and innovation program (grant agreement no. 853005), and the Honda Research Institute USA Inc.

## Notes

### Competing Interest Statement

The authors have declared no competing interest.

