## Supplementary Information for "Prediction of mycotoxin response of DNA-wrapped nanotube sensor with machine learning"

### Random Classification Model

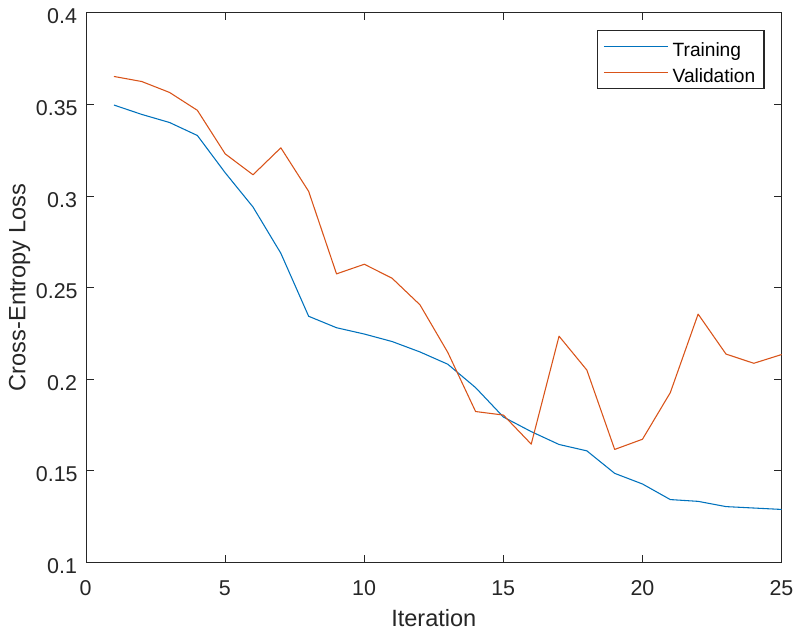

**Figure S1 –** Loss functions for 70 and 15% of data as train and validation in different stages of model classification: When the model is optimized using training data, the prediction is evaluated using validation data, and if the validation is not correctly predicted, the network will go back and adjust the model until the validation error is reduced (epoch of 14 was selected). Validation may also be utilized as a checkpoint at the conclusion of the training to avoid the ML training overfitting.

| 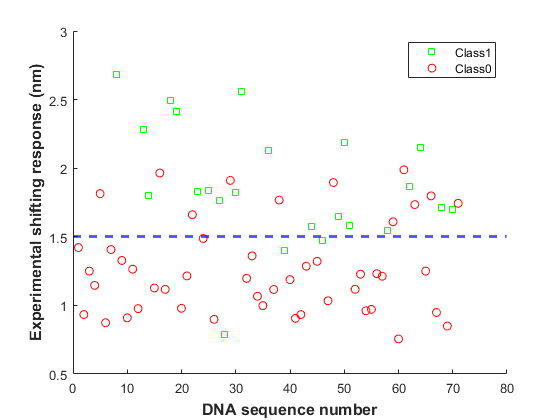  A | 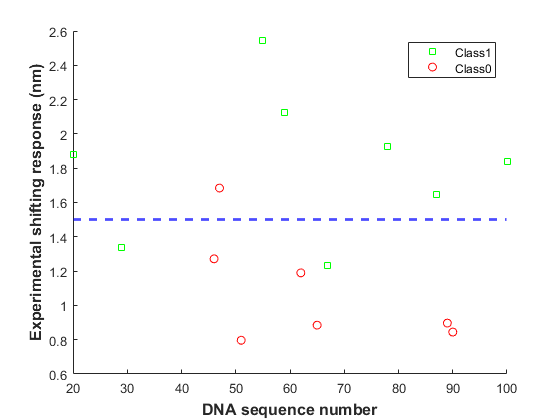 |
| --- | --- |
| 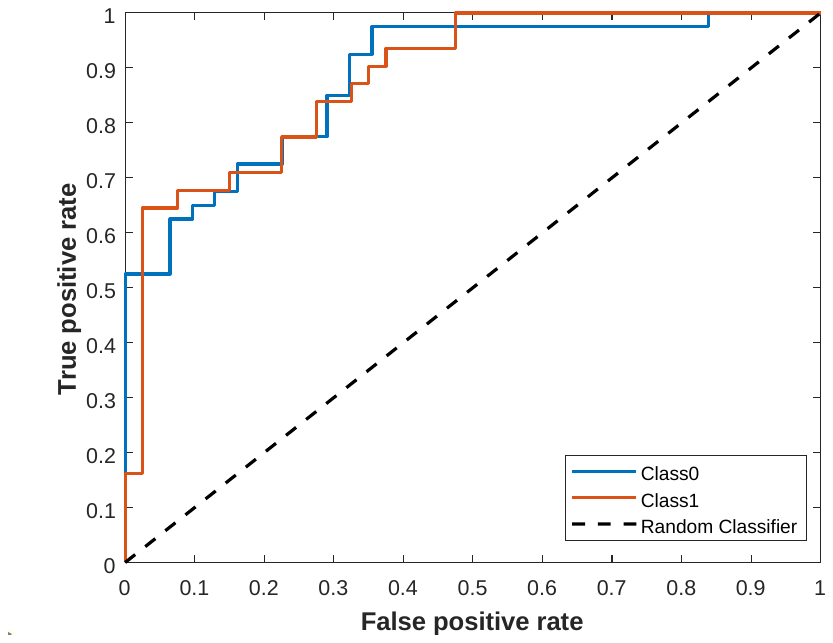  C | 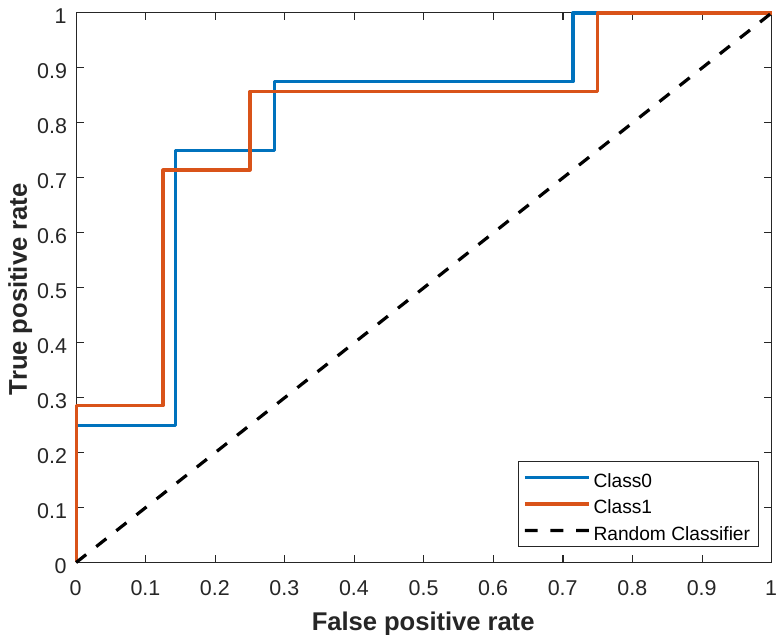 |

**Figure S2 – A and B)** Scatter plot for training, and validation data (Low shifting response (<1.5), Class 0 and high shifting response (≥1.5), Class 1) **C and D**) Receiver Operator Characteristic (ROC) curve for the training and validation data, classifiers that give curves closer to the top-left corner indicate better performance.

D

B

### Select the best classification model with cross-validation

**Table S1**. Evaluation parameters of ANN and SVM models with k-fold and one-leave-out cross-validations methods. The SVM model had more accurate results than ANN based on accuracy and AUC parameters.

| **Model/Cross Validation** | **Accuracy** | **AUC** |
| --- | --- | --- |
| **ANN - K-fold cross-validation** | 0.71 | 0.74 |
| **ANN - One leave out cross-validation** | 0.73 | 0.75 |
| **SVM - K-fold cross-validation** | 0.81 | 0.78 |
| **SVM - One leave out cross-validation** | 0.81 | 0.78 |

B

A

| 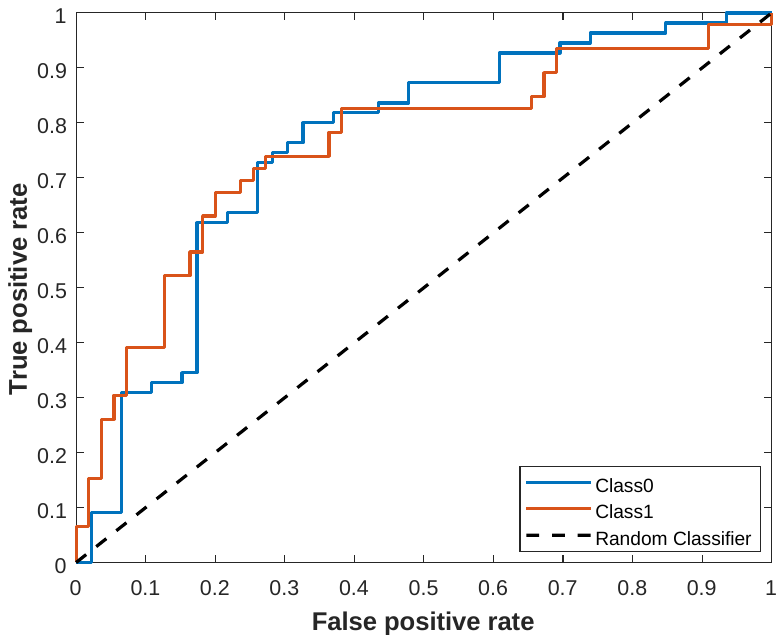 | 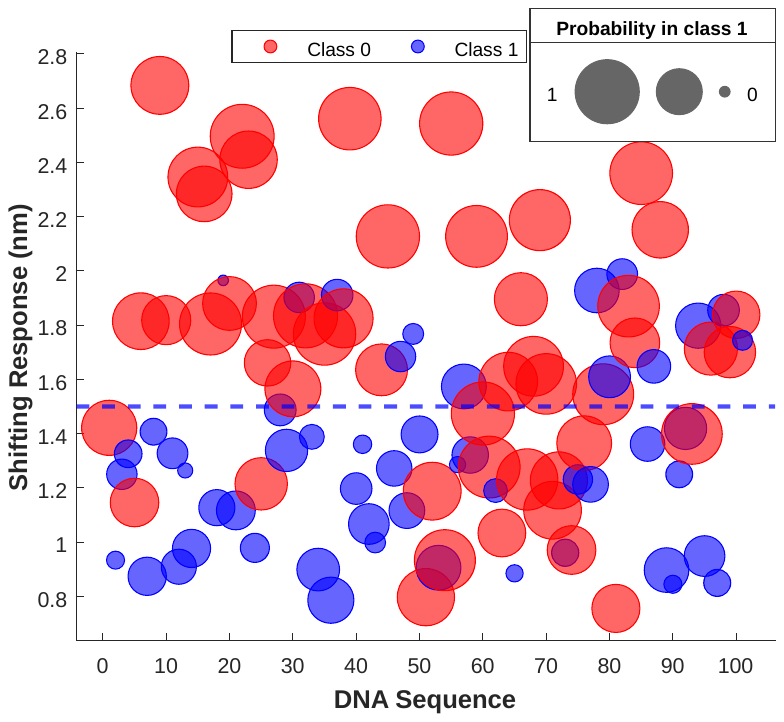 |
| --- | --- |

**Figure S3**-A and B) COF and ROC curve for ANN model with k-fold cross-validation: the AUC was measured at 0.71, C) The DNA sequence classification and probability of each DNA sequence (Size of the circle) which belongs to which class (Low shifting response (<1.5), Class 0 with red color and High shifting response (≥1.5), Class 1 with blue color) predicted by the k fold cross-validation and compared with experiment shifting response value to detection of mycotoxin by different DNA-SWCNT sensor.

**Table S2**- The result of the training data by different SVM method

| No. | Kernel function | Accuracy of K-fold  cross-validation |
| --- | --- | --- |
| 1 | Polynomial | 74 |
| 2 | Gaussian (Best Kernel) | 81 |
| 3 | Linear | 80 |
| 4 | Quadratic | 76 |

| 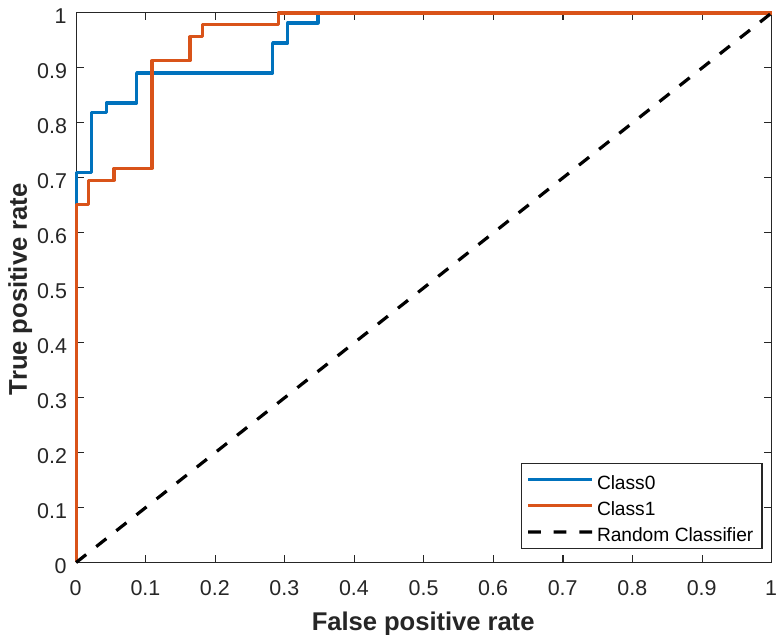  A | 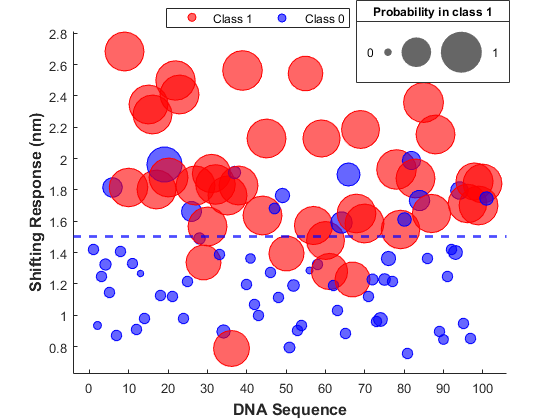  B |
| --- | --- |

**Figure S4**-A) The ROC curve for SVM model with k-fold cross-validation: the AUC was 0.7834, B) DNA sequences classification and presence probability of each DNA sequence in each class predicted by the SVM model and k-fold cross-validation compared with experimental shifting response values The size of the circles shows the probability values and the colors represent each class. In addition, low shifting responses (<1.5) show class 0 with blue color, and high shifting responses (≥1.5) show class 1 with red color.

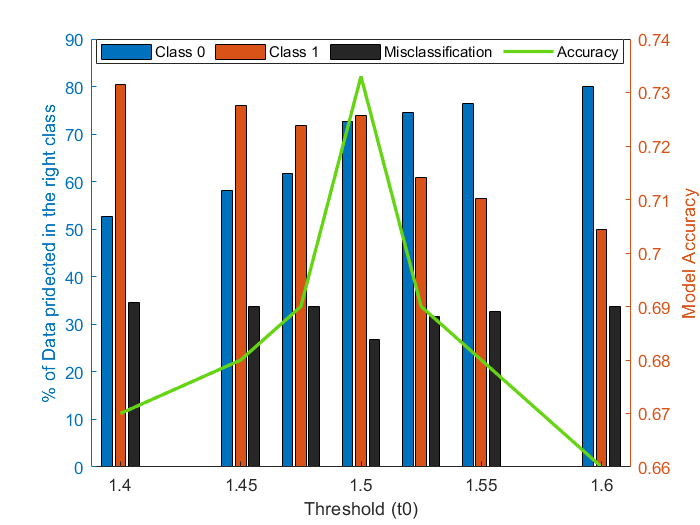

**Figure S5**- The effect of changing the threshold (t_0_) on the result of classification in 2 classes: The average value of t_0_ is 1.5 nm and the model was also trained with values higher and lower than average t_0_ by SVM model with one leave-out cross-validation. At average t_0_, almost all data is predicted in the right class with high model accuracy.

### Model’s ability to predict new dissimilar DNA Sequence

| 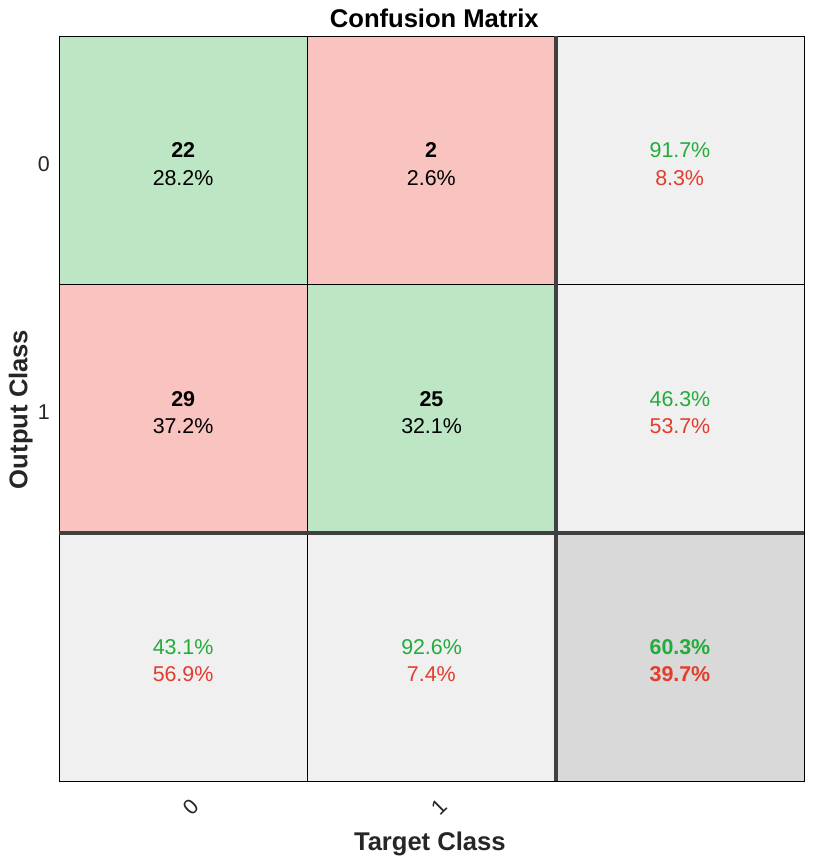  **A** | 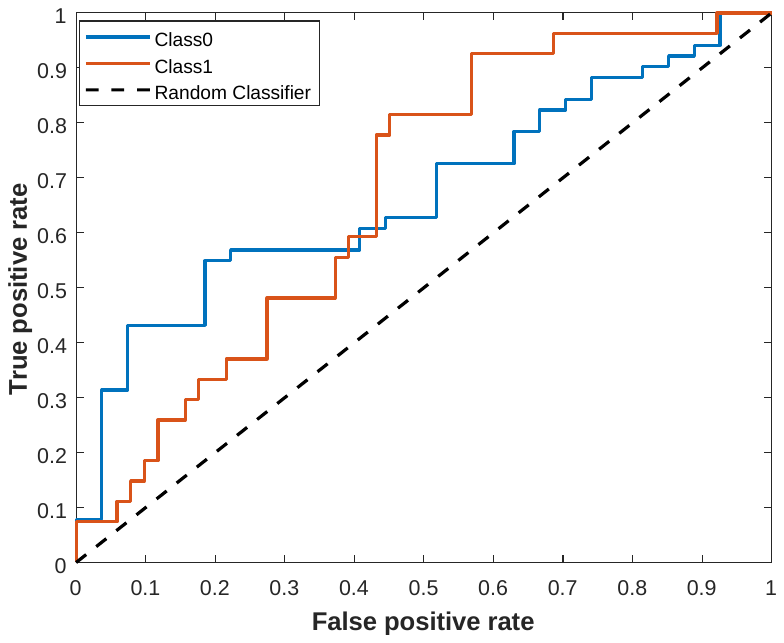 |
| --- | --- |
| 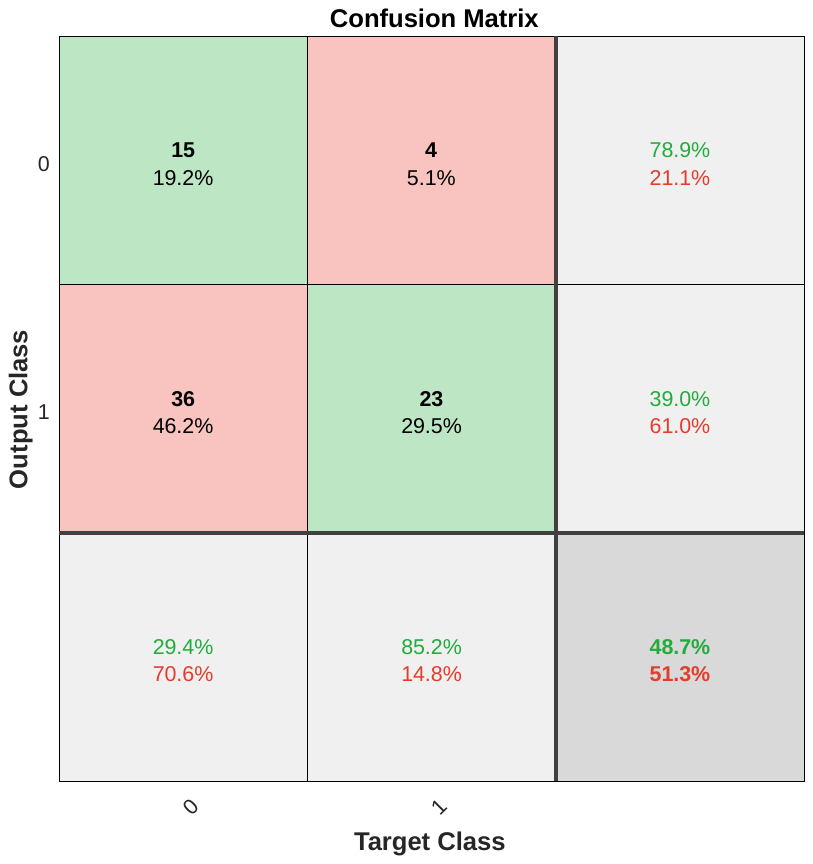  **B** | 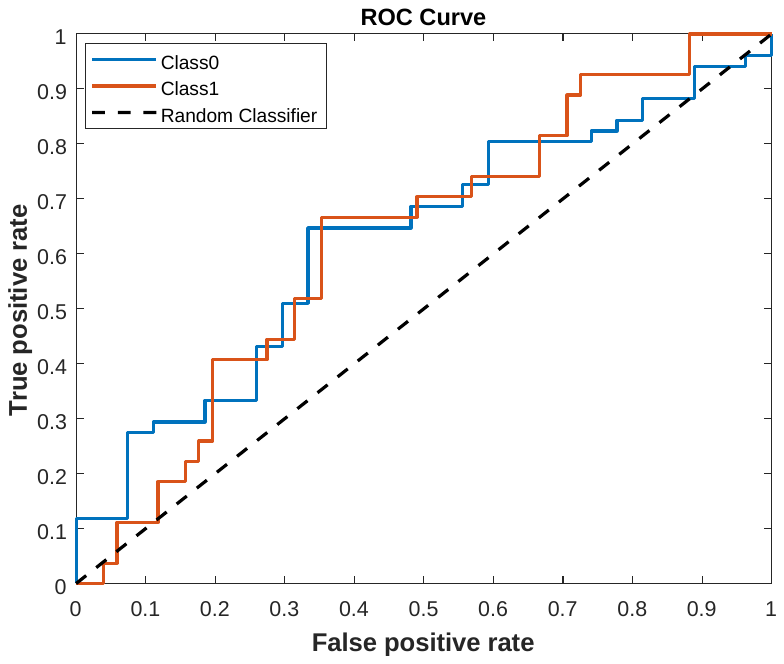 |

**Figure S6** –The COF and ROC curve for prediction of the new data by one leave-out cross-validation method for ANN model (A) and SVM model (B)

### 2.5 The ability of the model for finding new DNA Sequence in directed evolution method

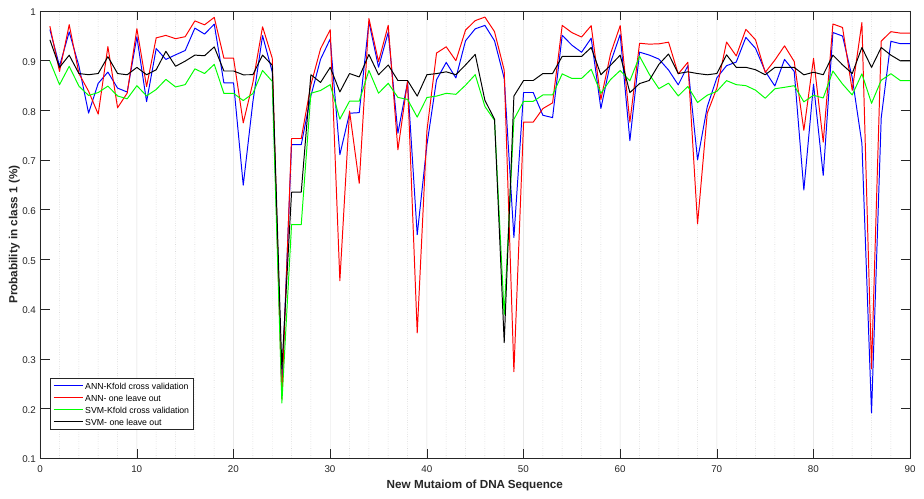

**Figure S7** - Prediction of new mutated DNA sequences by different ML models. The DNA sequence with the highest shifting response in the direct evolution cycle was selected as a parent sequence and then mutated by different 90 possibilities ((G…C)_30_ with a high shifting response of 2.7 nm)
